# Estimating gene conversion tract length and rate from PacBio HiFi data

**DOI:** 10.1101/2024.07.05.601865

**Authors:** Anders Poulsen Charmouh, Peter Porsborg Sørud, Lasse Thorup Hansen, Søren Besenbacher, Sofia Boeg Winge, Kristian Almstrup, Asger Hobolth, Thomas Bataillon, Mikkel Heide Schierup

## Abstract

Gene conversions are broadly defined as the transfer of genetic material from a ‘donor’ to an ‘acceptor’ sequence and can happen both in meiosis and mitosis. They are a subset of non-crossover events and, like crossover events, gene conversion can generate new combinations of alleles and counteract mutation load by reverting germline mutations through GC-biased gene conversion. Estimating gene conversion rate and the distribution of gene conversion tract lengths remains challenging. We present a new method for estimating tract length, rate and detection probability of non-crossover events directly in HiFi PacBio long read data. The method can be used to make inference from sequencing of gametes from a single individual. The method is unbiased even under low single nucleotide variant (SNV) densities and does not necessitate any demographic or evolutionary assumptions. We test the accuracy and robustness of our method using simulated datasets where we vary length of tracts, number of tracts, the genomic SNV density and levels of correlation between SNV density and NCO event position. Our simulations show that under low SNV densities, like those found in humans, only a minute fraction (∼2%) of NCO events are expected to become visible as gene conversions by moving at least one SNV. We finally illustrate our method by applying it to PacBio sequencing data from human sperm.

## Introduction

Recombination between chromosomes during meiosis is initiated by a double-strand break (DSB) of the chromosome, after which a single strand from the homologous chromosome can invade the broken double strand (strand invasion) to form a heteroduplex of DNA from different chromosomes (Jasin and Rothstein 2013). Depending on how the heteroduplex is resolved, strand invasion can result in either a crossover (CO) event or a non-crossover (NCO) event (Cole et al. 2010). A crossover event results in large segments of homologous chromosomes being swapped, whereas an NCO occurs when the DSB is repaired, using the homologous chromosome as a template, through the synthesis-dependent strand annealing pathway or the double Holliday junction pathway (Holliday 1964; Resnick 1976; Chen et al. 2007; McMahill et al. 2007). A subset of NCOs is observable as gene conversions, which occur when one or more SNVs are transferred non-reciprocally from one haplotype/chromosomal segment to another (Mansai et al. 2011; Lorenz and Mpaulo 2022). Gene conversions have several evolutionary consequences. For example, GC-biased gene conversion might serve to revert new germline mutations by opposing the AT mutation bias (Bengtsson 1985; Arbeithuber et al. 2015; Clessin et al. 2024), which has recently been shown to be potentially biasing the distribution of fitness effects of new mutations towards more beneficial mutations (Joseph 2024).

NCO events can be hard to detect. Firstly, an NCO event is only visible in genomic data as a new haplotype when it overlaps and converts at least one SNV from one donor haplotype to an acceptor haplotype. Because humans have low heterozygosity (approximately one site in 1500 are heterozygous), NCOs are hard to detect unless the number of bases affected by a single event is also more than 1000 bps. This seems to not be the case (Haldorsson et al. 2016; Arndt et al. 2023). Human NCO tract length, which is the length of double-stranded DNA converted in an NCO event, has been reported to be in the range 3-2086 bp, with a frequently cited estimate being 55-290 bp (Jeffreys & May 2004), making it less likely that any one NCO event will overlap and convert at least one SNV, hence becoming a (visible) gene conversion (Bosch et al. 2004; Jeffreys and May 2004; Hallast et al. 2013; Harpak et al. 2017; Hardarson et al. 2023). A related problem is that finding gene conversion events requires extensive sequencing if the gene conversion rate is low. Estimates of NCO rates in human span 5.9 · 10^−6^ to 8.75 · 10^−6^ NCO per base pair per generation, where this rate can be defined as the probability per generation that a base pair lies within an NCO tract (Williams et al. 2015; Halldorsson et al. 2016; Narasimhan et al. 2017). Furthermore, the sequence of both haplotypes needs to be known to determine which SNVs have been converted. Flanking unconverted SNVs on either side of the gene conversion event are typically necessary to distinguish a gene conversion event from a crossover event (Halldorsson et al. 2016; Hardarson et al. 2023).

Recombination events, including NCOs have often been studied by sequencing individuals in a pedigree spanning several generations or by using trios where the genome of mother, father and offspring is sequenced (Halldorsson et al. 2016; Porubsky et al. 2024; Prentout et al. 2024). While the idea of studying recombination events by sequencing gametes directly is old (sperm typing, e.g. Arnheim et al. 1991), recent studies are beginning to leverage new sequencing methods to directly study recombination events (Dréau et al. 2019; Carioscia & Weaver et al. 2022; Porsborg et al. 2024; Schweiger et al. 2024). The sequencing of highly accurate long reads from sperm samples offers a new way to identify gene conversion events. Specifically, if enough gametes from a single individual are sequenced, both parental haplotypes can be inferred such that gene conversions between two haplotypes at heterozygous sites can be directly observed (Porsborg et al. 2024; Schweiger et al. 2024). Furthermore, flanking SNVs on either side of the gene conversion event will typically be present such that the gene conversion event can be called. This is because HiFI PacBio long-read sequencing results in very long reads, usually with a mean read length above 10kb (Hon et al. 2020; Marx 2023) which greatly exceeds the expected size of conversion tracts.

Here, we build a model suitable for estimating NCO rate, tract length and detection probability directly from HiFi PacBio long read data (Figure 1). We use data simulations with varying levels of SNV density and correlations between SNV density and NCO intensity to study the statistical accuracy and robustness of our model. We then illustrate our approach with a recently obtained dataset (Porsborg et al 2024).

**Figure 1:**
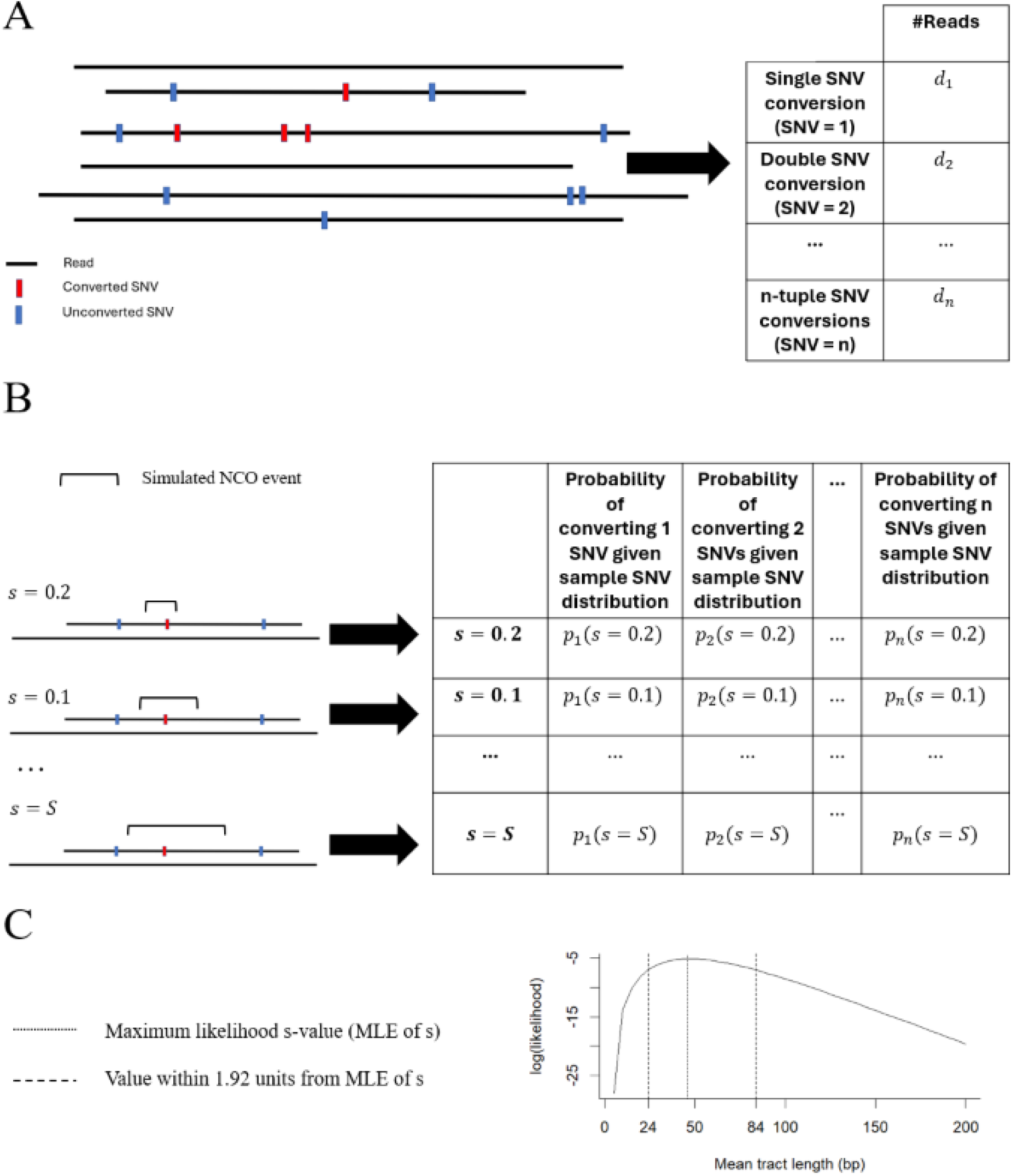
A method for estimating NCO tract length, rate and detection probability. A) The data. HiFi long reads were screened by Porsborg et al. (2024) for gene conversion, and read data is summarised via the counts of the number of reads where 1, 2,…, n SNVs are converted (conversion counts). B) The simulations. Using the SNV distribution along the sequenced sample genome, simulations are conducted with varying mean tract length and the probability of converting 1, 2,…,n SNVs (terms in eq. (5)) are estimated contingent on the SNV density and tract length distribution (conversion probabilities). C) The estimates. Using the conversion probabilities which take the non-uniform SNV distribution into account, the mean tract length which maximises the likelihood of the data (the conversion counts) is estimated using eq.(8).

## Results

### The Model

We count the number of gene conversion events where a single SNV has been converted and compare it to the number of gene conversion events where multiple SNVs have been converted. The ratio of these two counts provides information about the distributions of gene conversion tract lengths. This can be understood intuitively by realising that if single SNV conversions are far more common than multi-SNV conversion events, this indicates that tract lengths are typically quite short since it is unlikely for a short tract to convert multiple SNVs. This verbal expectation can be formalised as the expected probability of observing a single SNV conversion relative to the probability of observing a multi-SNV conversion.

We first derive this expected ratio of single to multi SNV conversions in the idealised case where all bases have the same probability of being SNVs (i.e. where the distribution of SNV positions is uniform along the genome). We then show that using the relative occurrence of single to multi-SNV conversions to estimate gene conversion tract length, rate, and detection probability can be generalised to any scenario wherein SNVs are clustered, as they are in real genomes. We do this by using simulations to obtain the conversion probabilities and then use the relative numbers of single and multi-SNV conversions in a maximum likelihood framework to obtain estimates for the mean tract length, rate and detection probability of gene conversion.

### The model – idealised case of uniform SNV distribution

Consider an NCO event with a tract length *L* base pairs (bp). For each bp, we assume that the probability of the gene tract terminating is *s* such that the probability of the gene conversion tract extending is 1 − *s*. Given this, the distribution of NCO tract lengths *L* follows a geometric distribution with parameter *s.* We initially assume a uniform SNV distribution such that for any base pair, the probability of observing a SNV is *p* and the probability of not observing a SNV is 1 − *p*. We initially assume that all SNVs overlapping the conversion event will be converted (this assumption can be relaxed, see Supporting Information S1).

We ask, given an NCO tract of length *m*, what is the probability, *P(#SNV=1 | L=*m), of converting a single SNV? In the idealised case where the SNV distribution is uniform (all positions in the tract have a probability *p* of containing an SNV), this probability is given by the binomial distribution,

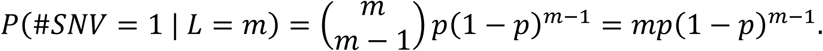

The joint probability of *L* = *m* and #*SNV* = 1 is then

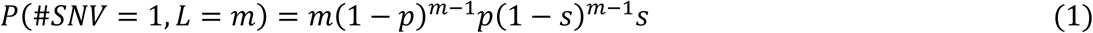

because tracts follow a geometric distribution, so we weight the probability of converting a single tract by the probability of sampling a tract of length *m* from a geometric distribution with mean 1/*s*.

Similarly, the joint probability of converting more than one SNVs given a tract of length *m*, *P(#SNV > 1, L=*m), is

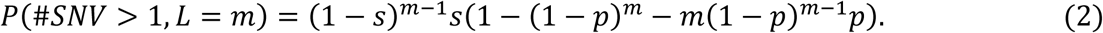

We now calculate P(#SNV=1) and P(#SNV > 1) by summing the probabilities over all possible values of *m.* This yields

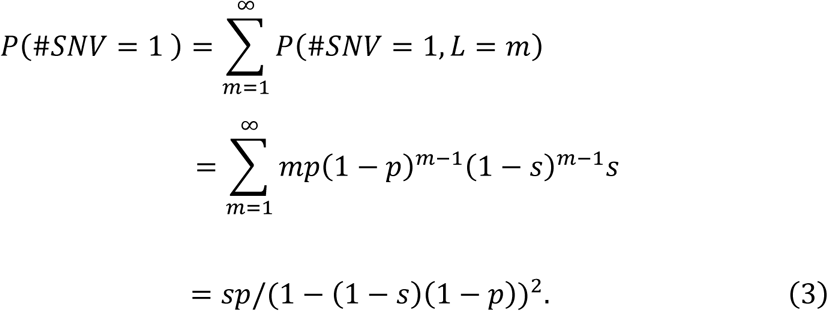

and

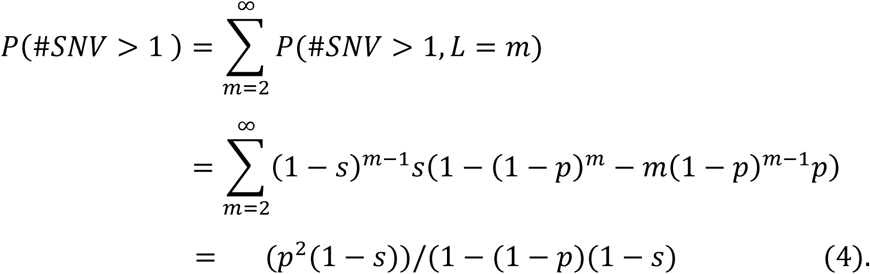

Notice that the sum in eq. (4) begins at *m=2*, because a tract length of at least 2 bp is necessary before the conversion of more than one SNV is possible.

The ratio of single to multi-SNV gene conversion events which we would expect to observe in the data, is now given as

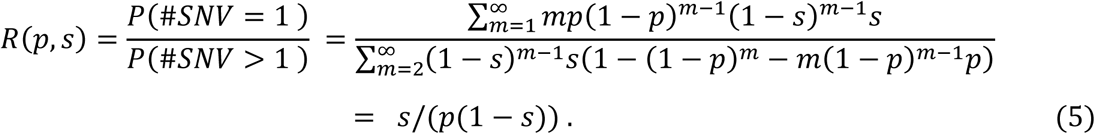

Notice that since the numerator and denominator of *R(p,s)* is the probability of converting one or more SNVs, respectively, we can also calculate the proportion of “silent” NCO events *S*(*p*, *s*), which we define as the proportion of NCOs converting no SNVs (i.e. the probability that an NCO does not become a gene conversion),

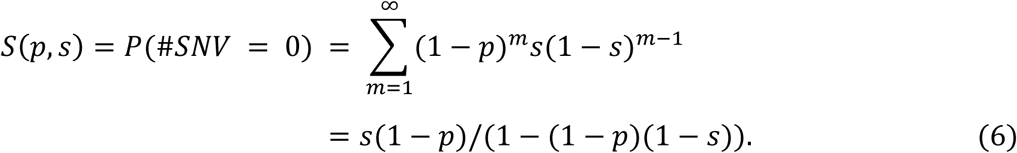

Since we know the proportion of NCOs which fail to convert a SNV relative to the proportion of NCOs resulting in observable gene conversions, we can now write an expression for the total number of expected NCOs in the whole genome D_E_, based on the observed number of gene conversions, D_o_

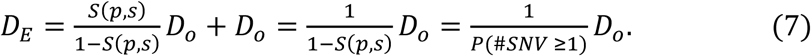

The total NCO rate, defined as the probability that a base is involved in an NCO event, is then simply *D*_*E*_ divided by the number of base pairs in the genome of interest, multiplied by the mean number of base pairs affected by an NCO event (i.e. the mean tract length). Note that to accurately estimate the NCO rate, S(p,s) was estimated by considering the distribution of reads length in our sample (see Supporting Information S2) for details.

### Estimating SNV conversion probabilities using the empirical SNV distribution

Different regions have very different coalescent histories. Times to the most recent common ancestor vary and accordingly modulate the SNV density, ranging from regions harbouring runs of homozygosity (recent coalescent event) to regions with a high density of SNVs (deep coalescent events) (Wiuf 2000b; Arndt et al. 2023). Additionally, variation in the mutation rate across the genome further contributes to a non-uniform SNV distribution (Oman et al. 2022; Barroso and Dutheil 2023). Because the assumption of SNVs placed uniformly is typically violated in genomes, and since this affects the accuracy of inference of the idealised model (see Supporting Information S3), we first estimated the probability of different conversion events (e.g. single, double, triple conversion etc) contingent on the exact SNV distribution. To estimate the probability of the different conversion events, we calculated (via simulation) the probability of converting 1, 2, …, n SNVs contingent on the SNV distribution in the genome of the sampled genome, using sequence data from Porsborg et al. (2024). We then calculated the proportion of different conversion events as a function of different values of *s* (see methods for details).

### Generalising to an arbitrary SNV distribution

Using a maximum likelihood framework, the idea that the relative numbers of single and multi-SNV conversion contain information about the tract length, rate and detection probability of gene conversion can be generalised to the case where SNVs are not uniformly distributed. We assume that a called set of gene conversion events is available from HiFi PacBio sequence data (e.g., Table 1). Out of these *k* gene conversion events, a certain number of events *d_1_, d_2_, …, d_n_* resulted in converting 1, 2,…, *n* SNVs, respectively (Fig 1A). We also assume that the distribution of SNVs in the genome of the sampled individual is known, the probabilities of converting 1, 2,…,n SNVs contingent on the SNV distribution (see methods) and given a mean tract length of 1/s (*P*_1_(*s*), *P*_2_(*s*), …, *P*_*n*_(*s*)) can be found (Figure 1B). Under these assumptions, the likelihood *L(s)* of *s* given the data *d_1_, d_2_, …, d_n_* can now be calculated as

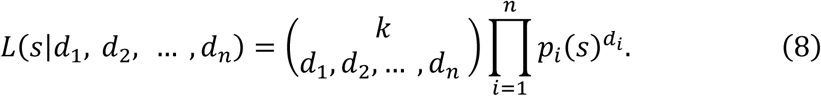

**Table 1:** Conversion counts obtained from a human sperm sample as reported in Porsborg et al. (2024) obtained by calling gene conversions in HiFi long reads. In this sample, 167 single SNV conversions, 12 double SNV conversions, 2 triple SNV conversion and 1 quadruple SNV conversion was observed.

|  | **1-SNV**<br>**conversion** | **2-SNV**<br>**conversion** | **3-SNV**<br>**conversion** | **4-SNV**<br>**conversion** |
| --- | --- | --- | --- | --- |
| **#reads** | 167 | 12 | 2 | 1 |

Finding the value *s*^∗^ which maximises *L(s)* yields a maximum likelihood estimate (MLE) of *s* given the data *d_1_, d_2_, …, d_n_* (Figure 1C).

We develop an analytical model which can be used to infer a parameter which describes the mean gene conversion tract length, the variance of gene conversion tract lengths, the NCO rate and the NCO detection probability based on the SNV density and the observed ratio of single to multi-SNV conversions (Figure 2). We tested the model by comparing analytical predictions of eq. (5) to simulations wherein gene conversion events were simulated across a genomic fragment of 30 Mb (see Methods for details). When the placement of a SNV position is uniform, we find that the analytical results are consistent with the simulation results, which means that the method allows for unbiased inference of gene conversion tract length, rate, and detection probability even when using data from populations with very low SNV density such as humans, i.e. populations wherein less than 1/1000 positions in the genome harbours heterozygous site (e.g. Zhao et al. 2003). This makes it suitable for inferring the mean NCO tract length for species with low levels of genetic diversity, such as humans (Figure 2).

**Figure 2:**
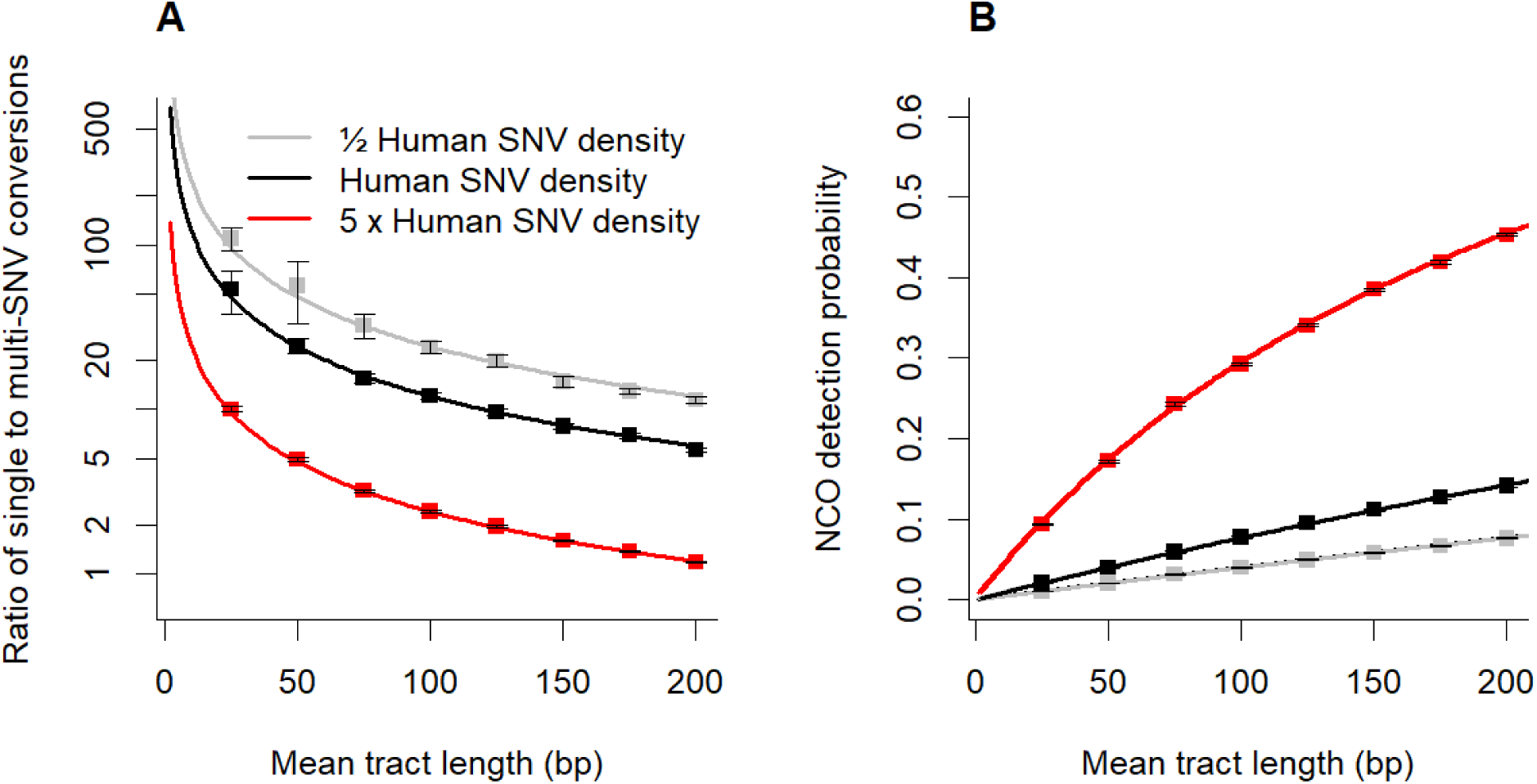
Analytical predictions of the ratio model (full lines) compared to simulation results (points denote means of 25 replicates) with 95% confidence intervals under different SNV densities, using approximately half the human SNV density as a proxy for the typical number of heterozygous sites (0.00083/2 SNVs/bp, e.g. Zhoa et al. 2003), human SNV density (0.00083 SNVs/bp) and five times the human SNV density (0.00083 · 5 SNVs/bp). The results show that in the idealised case where all positions in the genome have some probability of being an SNV, using the ratio of single to multi-SNV conversions can yield unbiased estimates despite very low SNV densities, such as those observed in populations of humans. (A). Eq. (5): Ratio of single to multi-SNV conversions as a function of mean tract length for three different SNV densities. (B) Detection probability as 1-S(p,s) (see eq.(6)) of all gene conversion events as a function of mean tract length for three different SNV densities.

In the HiFi long reads obtained from a human sperm sample (Porsborg et al. 2024), we observed a total of 182 gene conversions with between one and four SNVs converted per event (Table 1). Using the conversion counts, the maximum likelihood model presented (eq.8), the SNV distribution and the gene conversion events called by Porsborg et al. (2024), we estimate the mean tract length in a human sperm sample. We estimate the mean tract length to be 46 bp ([CI 95%: 24, 84], Figure 3A). In Porsborg et al. (2024) we present estimates from this and many additional samples using the method described here.

**Figure 3:**
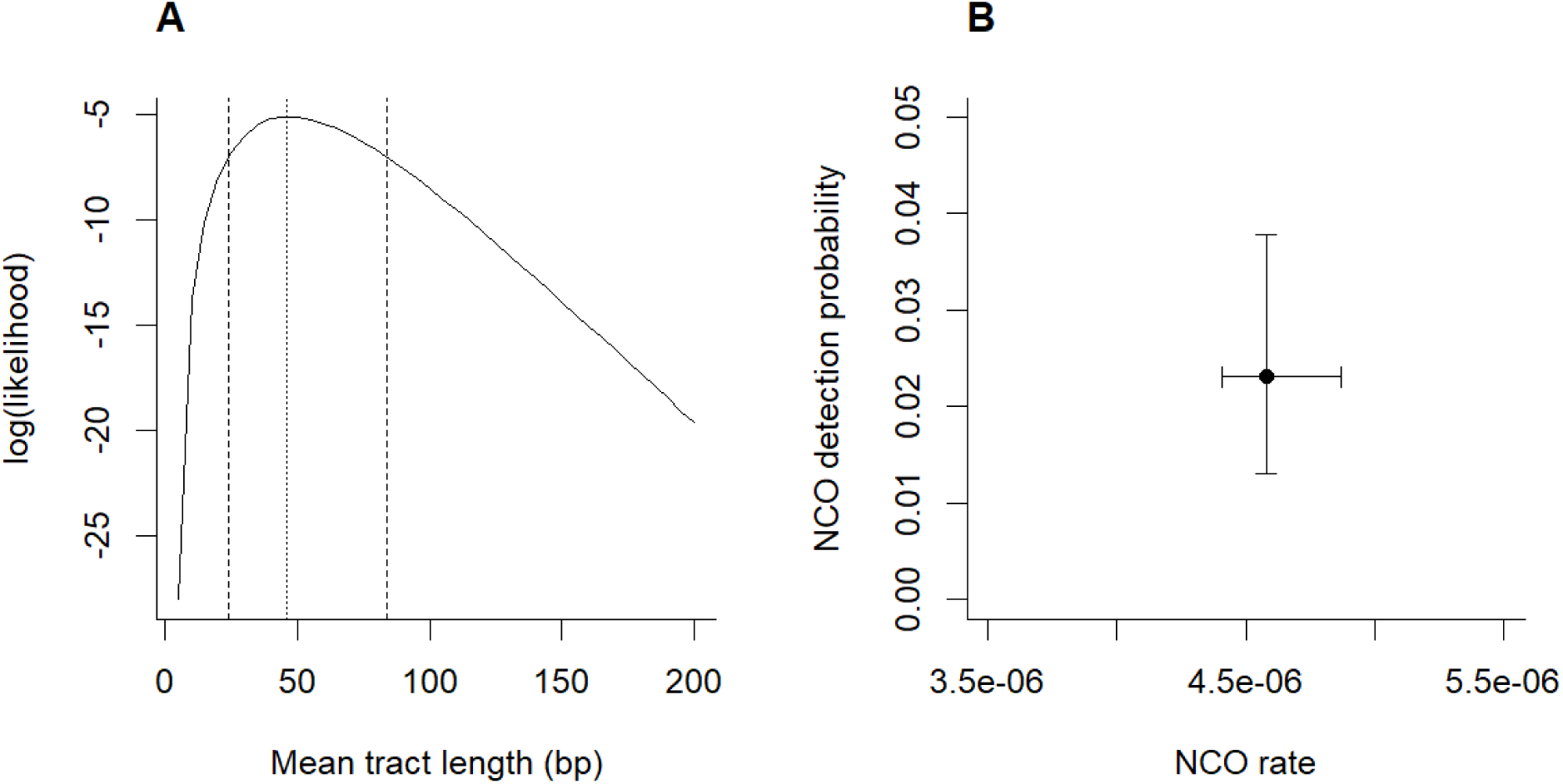
MLEs of mean gene conversion tract lengths, rate and detection probability inferred from gene conversion events called directly from HiFi PacBio data of sperm sample as obtained by Porsborg et al. (2024). A) Likelihood profile for mean tract length. Each point shows the log likelihood of the data (counts of single, double, triple,…,n-tuple gene conversion events for the sampled individual) conditional on the SNV distribution and density of the individual, i.e. eq. (8). Dotted vertical line represents the MLE and the vertical dashed lines show the 95% confidence interval. The results suggest that human gene conversion tracts are typically quite short (mean tract length of 46 bp). B). MLE of NCO detection probability (probability that an NCO becomes a gene conversion) and total NCO rate (including gene conversions) Bars denote the 95% confidence interval. The results indicate that most NCO events fail to convert at least one SNV meaning that these are not observable as gene conversions. Each MLE is based on 10^3^ simulations using the SNV distribution along the genome in the sample data.

We further find that given the MLE, the conversion counts (i.e. 167, 12, 2, 1) do not differ significantly (*p=0.71*) from the expected conversion counts when we assume tracts are sampled from a single geometric distribution parametrized with the MLE s (see Supporting Information S4).

The model also yields MLEs of the proportion of detected (and undetected) NCO events, that is, the proportion of NCO events which can be observed in data because they convert at least one SNV. We find that, on average, 2.31% of NCO events convert one or more SNVs, which means that most NCOs are ‘invisible’ because they happen to only convert identical sequences (Figure 3B). Dividing eq. (7) with the coverage and genome size, the MLE of *s* implies an NCO rate of 4.5810^-6^. Dividing eq. (7) with the coverage of the sample genome, we estimate that each male human gamete is expected to harbour, on average, 310 NCO events, of which approximately 7 are expected to become visible as gene conversions.

### Ascertainment bias and the effect of finite read lengths

We note that our pipeline for calling gene conversion events (Porsborg et al. 2024) requires that unconverted SNVs are present at the end of each read, such that it is possible to ascertain whether a read represents a potential crossover or gene conversion event. This leaves open the possibility of a slight ascertainment bias because some gene conversion events (e.g. those converting more than one SNV) might be indistinguishable from crossovers when occurring at the edge of a read. This ascertainment bias is very weak, since the average read length of HiFi PacBio reads typically greatly exceeds a typical NCO tract length. In the case of the data we analysed here, the average length of reads was 16.36 kb (see methods for details). We checked that simulating under the same read distribution as the sample (rather than assuming much longer reads) yields virtually the same probability of the different conversion events and, hence, almost the same MLEs (see Supporting Information S2).

### Robustness of MLEs under strong correlation between NCO positions and SNV density

The simulation approach to obtaining conversion probabilities assumes that all positions within the genome have the same probability of NCO events. However, if NCO events correlate (positively or negatively) with SNV density, our estimates of conversion probabilities could be inaccurate. We tested whether any positive or negative correlation between SNV density and NCO events existed by calculating SNV densities at double-strand breaks throughout the sample genome and comparing it to the overall SNV density. We found little to no correlation between double-strand breaks (DSB) (where NCO events are thought to be more likely to occur) and SNV density (CORR =-0.0012, P = 0.8225, see Supporting information S5).

While there is no evidence for a correlation between SNV density and NCO position in the data analysed here, this is not necessarily the case for all datasets. Furthermore, it is important to point out that the absence of correlation between DSBs and SNV density does not prove an absence of correlation between NCO and SNV density because a DSB can occur spontaneously (without being catalysed by recombination enzymes binding a DSB site) and then be repaired using sister chromatid. Because of this, we also tested the robustness of our model using simulations with different degrees of correlation between SNV density and NCO position. We found that strong positive or negative correlations between the position of NCO events and SNVs (see methods for details) result in only slight bias to the MLE obtained (Figure 4A). Specifically, if NCO positions show a strong positive correlation with SNVs, the MLE becomes slightly biased downwards since multi-SNV conversions become more probable, such that multi-SNV conversions (Table 1) can now be explained with a shorter mean tract length.

**Figure 4:**
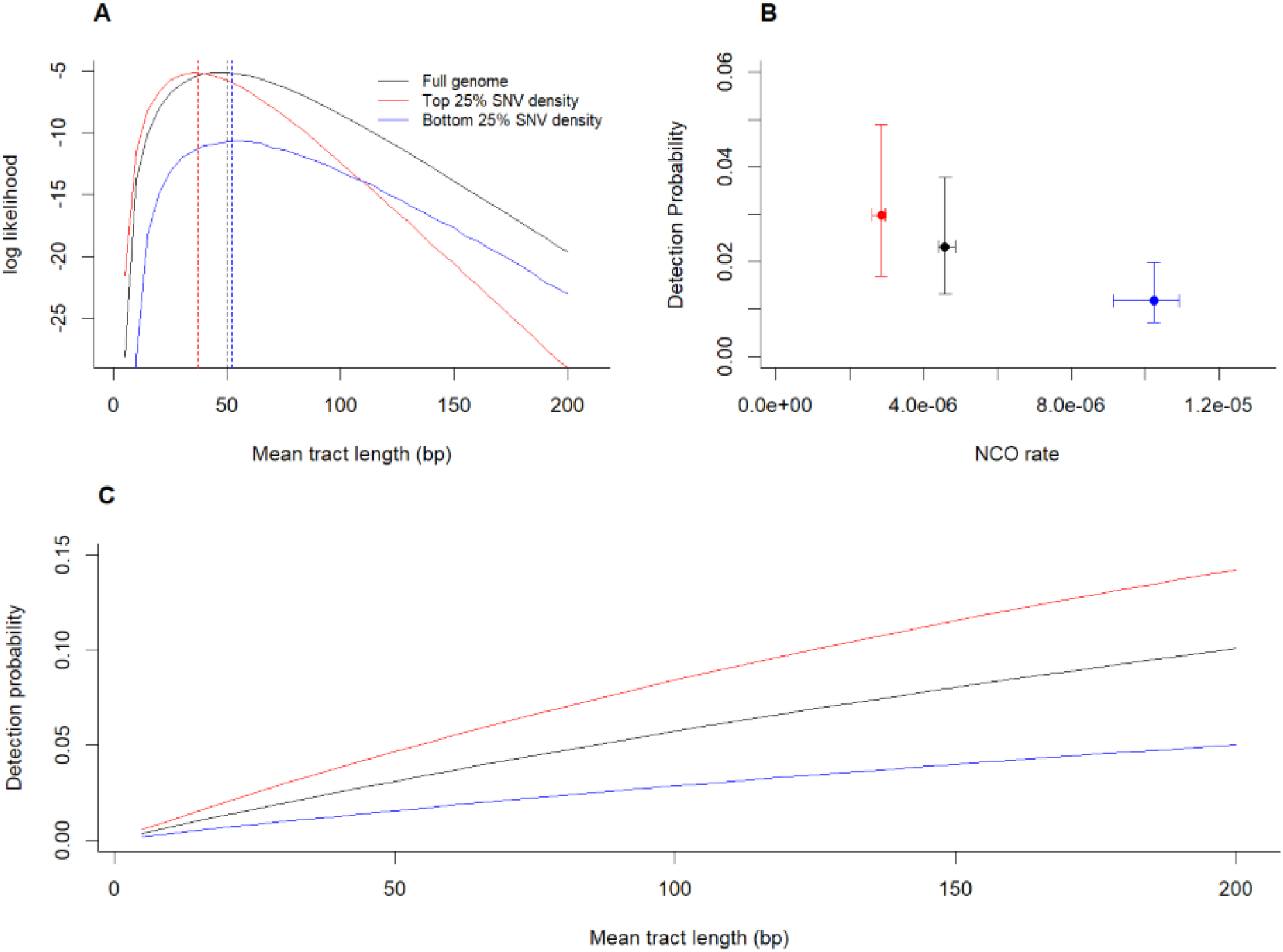
Comparison of MLEs under strong positive correlation between NCO position and SNV density (red), strong negative correlation between NCO position and SNV density (blue) and no correlation between NCO position and SNV density (black). (A) The MLE of tract length changes by −12 bp under strong positive correlation and +2 under strong negative correlation, suggesting the model is robust to correlation between NCO events and SNV density and overall heterogeneity of NCO positions along the genome. (B) Strong positive correlation between SNVs and NCO events can result in underestimation of the NCO rate whereas strong negative correlation can result in overestimation. (C.) Strong positive correlation between SNVs and NCOs can result in overestimation of the detection probability whereas strong negative correlation can result in underestimation. This is especially the case when tracts are long.

Conversely, a strong negative correlation between NCO position and SNVs results in a slight upward bias, since multi-SNV conversions become more unlikely, meaning that a longer mean tract length is required to explain the data (Table 1). Overall, we get MLE of tract length changing by up to −12bp under a strong positive correlation and +2 bp under a strong negative correlation. Specifically, compared to the case with no correlation between SNVs and NCO events, a strong positive correlation results in 26% underestimation, while a strong negative correlation results in a 4% overestimation of the mean tract length. This suggests that the model is robust even under strong positive or negative correlation, indicating that the model is applicable even when the assumption of a uniform distribution of NCO positions is strongly violated (Figure 4A).

Similarly, while strong positive or negative correlation between NCO events and SNVs affects the estimates of NCO rates (Figure 4B), these estimates (2.8610^-6^ and 1.0210^-5^) for strong positive and negative correlation, respectively) remain close to previous estimates (5.910^-6^ to 8.7510^-6^, e.g. Williams et al. 2015; Halldorsson et al. 2016; Narasimhan et al. 2017), and this tendency is reflected in the detection probabilities wherein negative correlation means fewer NCOs can be observed and positive correlations mean more NCOs can be observed as gene conversions (Figure 4B). This effect is amplified as tracts become larger (Figure 4C). Specifically, NCO detectability increases when NCOs only occur in the most SNV-rich regions of the genome, and NCO detectability decreases when NCOs only occur in regions with low diversity. This also means that a negative correlation results in a higher estimate of the NCO rate since a low detectability implies many unseen NCOs.

Conversely, positive correlation reduces the estimated NCO rate since the detectability is increased. Detection probability increases when NCOs occur in SNV-rich regions since more NCOs will become gene conversions as the chance of converting at least one SNV increases. Similarly, detection probability decreases when NCOs occur in low-diversity regions since fewer NCOs will convert at least one SNV. Despite the MLE being robust to correlation, it is important to point out that the correlation between SNV density and NCO events has a much greater effect on NCO rate estimates than NCO tract length estimates. The same applies to NCO detection probability, where positive or negative correlation can result in a ∼2-fold difference in the estimated detection probability.

We also investigated the accuracy of inference of the idealised model (under uniform SNV distribution) and the maximum likelihood model (under the SNV distribution of the sample genome). Both models were tested under varying tract lengths and varying amounts of data i.e. observed gene conversion tracts (for details, see methods). We found that both models are unbiased as the estimated mean coincided with the true mean across tract lengths (Figure 5). The sampling variance of estimates increases under both models when the sample size (i.e. the number of observed tracts) is low, and when the average tract length is short (cf. Figure 5A-B, Figure. 5E-F). At low mean tract length, individual estimates vary more because changes in the ratio have a large effect on mean tract length when tracts are short (Fig S2.2). However, when NCO tracts are short, the variance in the number of single to multi-SNV tracts is also expected to be the lowest (this is because the variance of the binomial distribution is *nq*(1 − *q*) where *q* is the probability of a single SNV tract, and *n* is the number of observed tracts). This former effect seems to be dampened by the latter, such that even when the true mean tract length is short (such as we estimate in this study, i.e. a mean tract length ∼50 bp), the estimated mean coincides with the true mean and the majority of all individual estimates are with +/− 25 bp of the true tract length, even with nearly 4 times less data that used in the present study (Figure 5). This suggests accurate estimates can be obtained even when coverage (and hence the number of observed tracts) is 3-4 times lower than in our data example (mean coverage of 26; see methods for details). The same is true for the idealised model (Figure 5; See also Supporting Information S6).

**Figure 5:**
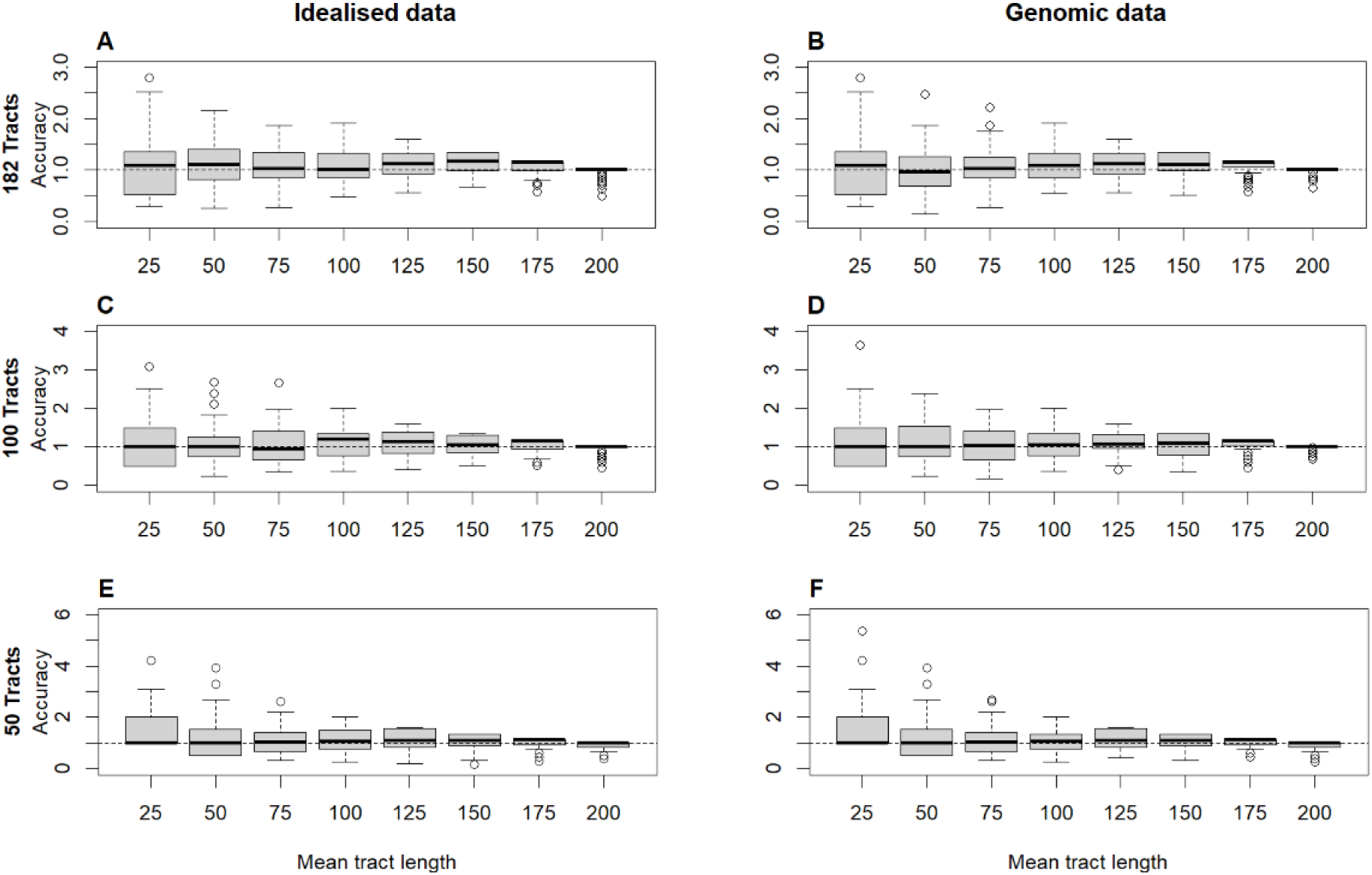
Accuracy of inference under (A, C, E) the idealised model (uniform SNV distribution) and (B, D, F) the maximum likelihood model (used to perform inference genomic data). Accuracy of inference is shown under different sample sizes (i.e. different number of identified tracts): 182 tracts, as used in this study (A-B), 100 tracts (C-D), and 50 tracts (E-F). Dashed line show perfect accuracy (estimate tract length / true tract length) = 1. Each boxplot contains 100 replicates (see Methods for details).

## Discussion

We developed a method for estimating the rate and length distribution of NCO events. We showed that this method is unbiased even for organisms with low SNV densities, such as humans, and that sufficient input data for the method can be obtained from sequencing gametes from a single individual and then calling gene conversion events directly from HiFi PacBio sequencing data.

We apply our method to a data set from humans (Porsborg et al. 2024) which estimates consistent with previous studies. While the dataset analysed here is limited to a single individual, and inter-person variation in gene conversion tract length and rate is certainly possible, we note that the confidence intervals of the estimate we obtain overlap with the confidence intervals of several previous estimates obtained from human data (Jeffreys & May 2004; Halldorsson et al. 2023, Schweiger et al. 2024. See also Supporting information S7).

While we use HiFi PacBio data obtained by sperm sequencing for illustration here, inference of NCO rate, tract length and detection probability could easily be done with a different type of data and different organisms – all that is required is the number of gene conversion events counts (i.e. counts of NCO events resulting in single, double, triple, etc. SNVs conversions) and the background SNV distribution of the sample where NCOs were called. Because of this, our method could also be used to obtain estimates by using gene conversions called in trio data (Halldorsson et al. 2016), in hybrid cross experiments (Li et al. 2019) and in pollen or sperm-typing (Lien et al. 2000; Jeffreys and May 2004; Tiemann-Boege et al. 2006). However, comparing estimates obtained from very different data types is not necessarily straightforward since different ways of obtaining data can lead to different ascertainment biases of NCO events. For example, trio data makes it in principle possible to observe NCOs inducing gene conversion tracts of an arbitrary length whereas the finite read length constrains HiFi PacBio long read data. On the other hand, when all individuals within a trio are heterozygous at a specific position, phasing may be incomplete (Miller and Piccolo 2021). A more detailed investigation of the extent to which different types of datasets might bias the estimates of NCO rate and tract lengths upwards or downwards would be a worthwhile aim for future work.

We exploit the fact that multi-SNV conversion becomes more common as the tract length increases, a finding that has also been made in early simulation studies (Gay et al. 2007). More recently, Li et al. (2019) proposed a method which assumes an exponential distribution of gene conversion tracts and uses information about the co-conversion of SNVs. Rather than using the ratio of single to multi-marker conversion events, the method of Li et al. (2019) creates a composite likelihood function by taking all consecutive pairs of SNVs adjacent to or within gene conversion tracts and considering whether these were part of the same conversion tract or not, as a function of the distance between the SNVs. While the method also takes the distance between potentially co-converted SNVs into account, it results in a composite likelihood (pseudolikelihood) because SNVs within the same conversion tract are assumed to be independent, which is not the case. Considering the ratio of single to all multi-SNV events (as in our model) has the advantage that it grants analytical solutions for the probability of an unobserved NCO (i.e. ‘invisible’ gene conversion) and the total NCO rate in the idealised case. Combining these methods, such that the distances between converted markers are also considered, could result in more precise predictions. Exploring this could be fertile grounds for future work.

### Gene conversion tract length and the geometric distribution

Our method assumes that tract length follows a geometric distribution. We consider this assumption to be reasonable since several studies have found the geometric distribution (or the continuous version of the distribution, the exponential distribution) to fit well with data (Hilliker et al. 1994; Taghian and Nickoloff 1997; Li et al. 2019) and this was also the case here when considering the conversion counts (see supporting results S4). This makes sense because the transfer of SNVs from one sequence to another via gene conversion is, at the core, the result of a polymerase moving along a sequence, which is being repaired using some donor sequence as the template (Jasin and Rothstein 2013). If the polymerase has a certain probability of stopping at each base and the remaining probability of extending the tract at each base, this will result in a geometric distribution of tract lengths (Wiuf 2000a; Frisse et al. 2001; Padhukasahasram and Rannala 2011; Setter et al. 2022). A recent study found that a mixture of two negative binomial distributions fitted datasets on gene conversion tract lengths better than a geometric distribution (Hardarson et al. 2023). We have compared estimates from different studies and methods directly on the same data and commented on some relevant differences (Supporting Information S7).

In some studies, a small subset of the observed gene conversion tracts seems incompatible with viewing all tracts as coming from a single geometric distribution with a low mean. Several recent studies have pointed out that while the majority of gene conversion tracts appear to fit a geometric distribution, the distribution of gene conversion tracts as a whole does not, since a very small fraction of tracts are much larger than expected under a single geometric distribution (Halldorsson et al. 2016; Wall et al. 2022; Versoza et al. 2023). In a study of olive baboons (*Papio anubis*), Wall et al. (2022) found >99% of all gene conversion tracts to be short, whereas a few tracts were very long (mean length 47.58kb). Similar results have been found in a study of rhesus macaques (*Macaca mulatta*) where two very long tracts were observed, the mean length of which was 5.166 kb (Versoza et al. 2023). In human data, Halldorsson et al. (2016) also reported a minority of very long NCO tracts, the mode of which was 30 kb.

Since the distribution of gene conversion tract lengths appears to be bimodal, this suggests (as proposed by Hardorsson et al. 2023) that two distinct processes might give rise to gene conversions – one which is responsible for nearly all tract and results in short tracts, and one which is very rare, but results in long tracts. While fitting a mixture distribution improves model likelihood (Hardarsson et al. 2023; Schweiger et al. 2024), doing so requires far more data since long tracts are rare. Specifically, the current dataset from Porsborg et al. (2024) comprises 182 tracts, but if ∼1% of all tracts are long, a sample of 182 tracts would likely only include 1-2 long tracts, which is insufficient information to describe a whole distribution. Thus, while a mixture distribution is likely a more accurate representation of all NCO events, a mixture model requires far more data and, given the additional complexity of the model, may result in wider confidence intervals around estimates. When using HiFi PacBio long read data, these rare long NCO events can, in principle, be detected if the distribution of read lengths overlaps with the distribution of long NCO event lengths (see Supporting Information S8). Since the short NCO events comprise both the majority of all NCO events and the majority of all converted SNVs (e.g. Wall et al. 2022; Schweiger et al. 2024), we believe model fitting single distribution is useful, although given additional data, the approach we described here can easily be extended to a mixture distribution (see Appendix; see also Supporting Information 9).

## Methods

### Simulation set-up, initialization, and parameters

The likelihood function requires conversion probabilities contingent on the SNV distribution in order to estimate the mean tract length, NCO rate, and NCO detection probability. We estimate these conversion probabilities by simulation. In each simulation, NCOs occurred at 10^5^ positions throughout the genome of the sample. These positions were chosen from a uniform distribution *U*(1, *N*) where *N* is the genome size (we also assess the robustness of the model when the assumption of a uniform distribution of NCO positions is violated. We find that strong correlation causes only slight changes to the MLE – see results for details). The tract length of the gene conversion event was sampled from a geometric distribution with mean 1/*s*. The proportion of NCOs converting one, two, *n* SNV under each value of *s* (hereafter denoted *P*_1_(*s*), *P*_2_(*s*), …, *P*_*n*_(*s*)) was calculated. All classes up to *n=11* were calculated for each simulation. Integers in the range [5, 200] were used for *s* in the simulations. For each value of *s,* we simulated the entire genome 10^3^ times (i.e 10^3^ simulation replicates) and mean values of *P*_1_(*s*), *P*_2_(*s*), …, *P*_*n*_(*s*) were calculated for each *s*. We note that when the SNV distribution is non-uniform, eq. (6) cannot be used to estimate NCO detection probability. Because of this, we the simulation output to estimate NCO detection probability as 1 − *P*_0_(*s*) since by definition, this is the proportion of NCOs moving at least one SNV, hence being detectable.

Simulations were conducted on a slurm cluster (slurm 23.02.5) and the full source code is available at [https://github.com/r02ap19/GeneConv] along with a ReadMe file detailing how to change model parameters as well as how to compile and run the code.

### Testing model robustness under strong heterogeneity in recombination position

Recombination events, including NCO events, are thought to have a higher probability of occurring in recombination hotspots. If the positions of NCO are not random with respect to SNV density, the conversion probabilities estimated under the assumption of a uniform distribution of NCO positions, and hence the resulting MLE, could be inaccurate. We tested the robustness of the MLE obtained when the position of NCO correlated strongly (negatively or positively) with the SNV density. This was done by counting all SNV in 1 Mb windows of the sample genome. Conversions probabilities and MLE estimates where then obtained by simulation of data in two different cases: one where NCOs only occurred randomly within in the top 25% most SNV rich 1 Mb windows of the genome (strongly positive correlation between NCO position and SNV density) and one where all NCO events only occurred randomly within the bottom 25% least SNV rich 1 MB windows of the genome.

We further quantified any positive or negative correlation between SNV density and NCO position by calculating SNV density around all previously detected double-strand breaks found in the sample genome (Pratto et al. 2014) compared to the SNV density at random 20kb positions across the genome (for details, see Supporting Information S5).

### Accuracy of inference

We tested the accuracy of inference of the idealised model by running 100 simulation replicates of a 30 Mb genome fragment with the human SNV density of 0.00083 (e.g. Zhao et al. 2003) which was exposed to 1000 NCO events at random positions. Note that the chosen size of the genomic fragment and number of NCO events does not affect accuracy as long as the genome is sufficiently large to make overlap between NCO events unlikely (as in real genomes). For more details on the minimum number of observed single/multi-conversion events necessary to use the idealised model, see Supporting Information S6. The ratio of single to multi-SNV conversion was then calculated, and eq. (5) was then solved for an estimate of the mean tract length. This was done for a range of tract lengths in the interval [25, 200] (Figure 5).

We also tested the accuracy of inference of the maximum likelihood model. Here, we calculated (by simulation) the probabilities of converting 1, 2,…, n SNVs (conversion probabilities) contingent on the SNV distribution of the sample and given a set of mean tract lengths in the interval [25, 200]. We then sampled tracts (182, 100, or 50) continuing 1, 2,…, n SNVs relative to the conversion probabilities. Given the sampled tracts, the mean tract length which maximised eq. (8) was found, and this was the estimated mean tract length. For both models, this was replicated 100 times for each tract length. For both the maximum likelihood model and the idealised model, accuracy was defined as estimate/true value, hence accuracy = 1 denotes perfect accuracy (Figure 5).

### Calling gene conversions in HiFi PacBio data

We used data from Porsborg et al. (2024) which describes a pipeline comprising sample preparation, sequencing, reads filtering based on quality, calling of gene conversion candidate reads, and curation of candidate gene conversion reads (Porsborg et al. 2024). In the following, we give a brief outline of their pipeline.

One human sperm sample (HS25, see Porsborg et al. 2024) was obtained from an approximately 25 year old anonymous man (because the donor is anonymous, the exact age is unknown). Purified sperm were used because gametes are thought to undergo more gene conversion than somatic cells (Porsborg et al. 2024). The sperm from the ejaculate was purified using a density gradient centrifugation and sequenced at 26X mean coverage using PacbioHiFI sequencing. This results in a set of consensus reads originating from different sperm cells present in the ejaculate. The average read length of the consensus reads were 16.36 Kb, and these were used to create a high-quality de novo genome assembly which spanned >95% of the genome with N50 contig size of 70.7 and with 97.9% of all contigs being larger than 1Mb. The consensus reads were then mapped back to the de novo assembly and all high-confidence SNVs were called while SNVs were assigned to haplotypes resulting in full phasing of all variants. This allowed for gene conversion events to be called as a switch back and forth between the two haplotypes on a single read, stemming from the fact that at least one SNV was transferred unidirectionally from one haplotype to another (see Porsborg et al. 2024 for more details. See also Schweiger et al. 2024 for a similar approach).

A total of 200 gene conversion candidate reads were called and manually curated using IGV (Robinson et al. 2011) to remove likely false positives resulting from large indels or mapping errors. 182 of the candidate gene conversion events were approved following manual curation and these events were used to obtain estimates of gene conversion tract length, rate and detect probability. The approved events resulted in the transfer of between one and four SNVs (Table 1). A full description of this pipeline, along with all the relevant scripts is given by Porsborg et al. (2024). We note that enough gene conversion events to obtain accurate estimates can be called with far fewer events than 182, which means that 2-3 times lower coverage would likely have yielded the same accuracy (see Results). Further, using specifically HiFi PacBio data is not strictly necessary since, our method can be applied to any dataset wherein SNVs and gene conversion events have been identified and the number of single and multi-SNV transfers has been counted.

## Acknowledgements

We would like to thank Regev Schweiger and Richard Durbin for their insightful comments. Some of the computing for this project was performed on the GenomeDK cluster. We thank GenomeDK and Aarhus University for providing computational resources and support that contributed to these research results. We also thank The Novo Nordisk Foundation, The Independent Research Fund Denmark and The European Research Council for funding (for details, see Funding).

## Author contributions

MHS, TB and APC conceived the study. APC wrote the manuscript with substantial input from MHS, TB, AH, SB and LTH. APC developed the model with substantial input from AH, TB and MHS. APC wrote the simulation software. PSP called SNVs and GC events and curated the HiFi long-read data. APC analysed the data using the simulation-based framework. KA and SBW were responsible for all lab-related handling and processing of the sperm sample. LTH wrote the Appendix with substantial input from AH.

## Data availability statement

All simulation software is available at [https://github.com/r02ap19/GeneConv] along with a readme file describing how to compile and run the code. The repository also contains the input data file. Gene conversion counts were obtained from Porsborg et al. 2024 (BIORXIV/2024/601967).

