## Supporting Information for "Estimating gene conversion tract length and rate from PacBio HiFi data"

**S1: Model accuracy under incomplete penetrance of NCO events…….………………....2**

**S2: Read length effects on single/multi-SNV conversion calls and detection………..…...4**

**S3: The necessity of accounting for non-uniform SNV distributions……………………..5**

**S4: Model fit to gene conversion counts…………………………………….…………..…..7**

**S5: SNV density around double-strand breaks………………………………………….....7**

**S6: Inference ability given sample size (idealised SNV distribution)………………...…...9**

**S7: Comparison of MLE to previous studies and methods……………………...…….…10**

**S8: Detectability of long NCO events in HiFi long reads data…………...…..…………..13**

**S9: Direct comparison between simulation procedure and recursion method…….……14**

**S1– Model accuracy under incomplete penetrance of NCO events**

Following Hardarson et al. (2023), we define penetrance as the probability that a SNV within an NCO tract is converted. That is, under complete penetrance (P=1), all SNVs within an NCO tract will become visible as gene conversions whereas under incomplete penetrance (P< 1), SNVs within an NCO tract have the probability P of becoming gene conversions.

To quantify the effect of incomplete penetrance, we tested the accuracy of the idealised model under a uniform SNV distribution by inferring mean tract length from ratios obtained from simulation with different levels of penetrance (Figure S1.1A). We also tested the accuracy of the model when this assumption was relaxed by seeing whether the MLE of the mean tract length would change under different levels of penetrance when inferring a mean tract length given the gene conversion counts and SNV distribution of one sample (Figure S1.1A). In accordance with Hardarson et al. (2023), our results show that assuming full penetrance in cases where the penetrance is incomplete (i.e. below 1) results in a downwards bias in estimation. Here, we have used a range of values for P chosen to span the estimates of P obtained by Hardarson et al. (2023) by applying their likelihood model to data from mice (Li et al 2019) and data from humans (Halldorsson et al. (2016)).

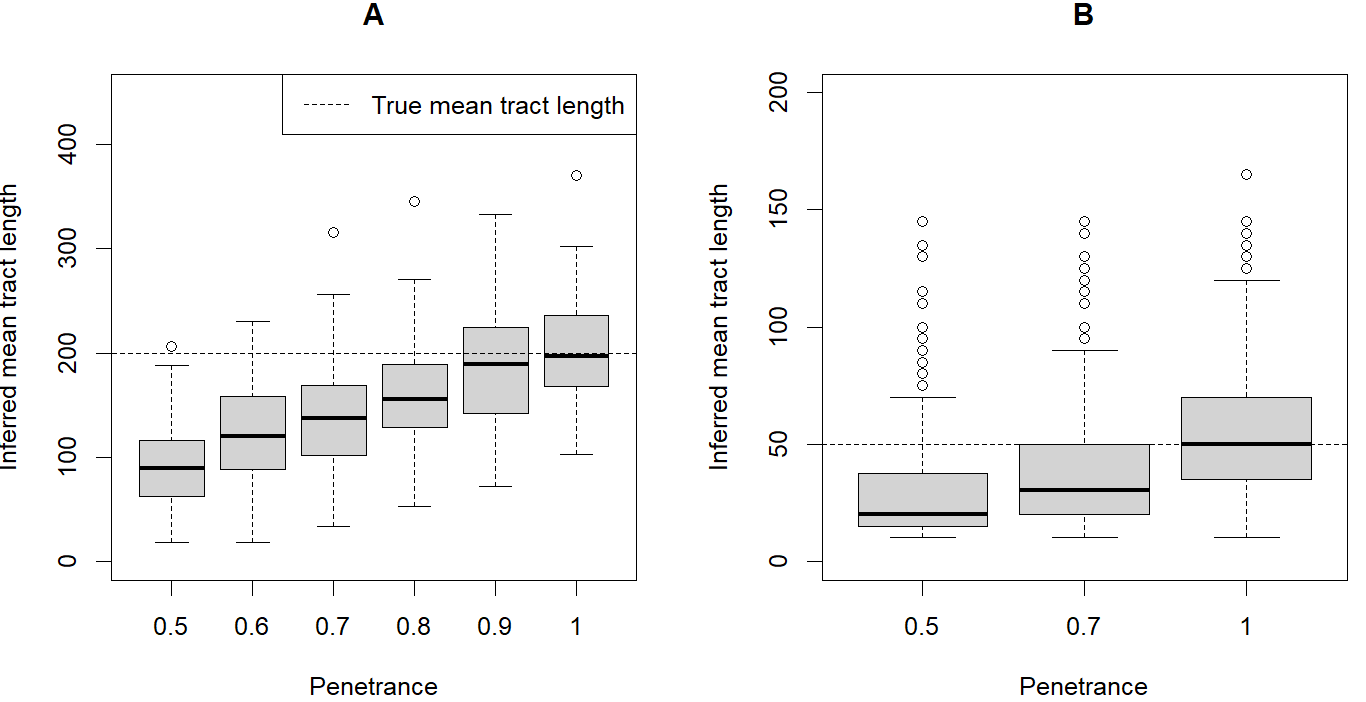

*Figure S1.1: The accuracy of inference under different levels of penetrance for the idealised case of uniform SNV distribution (A) and given the SNV distribution of our sampled individual (B). For the idealised case, estimates in boxplots were created by solving eq. (5) (in the main text) for ratios obtained from simulations. For the data case, estimates in boxplots were created by finding the s which maximised the likelihood (eq.(8) in the main text) given the gene conversion counts (see table 1 in the main text). Each boxplot is based on 100 simulations. The penetrance values cover those explored in Hardarson et al. (2023). The results show that, as pointed out by Hardarson et al. (2023), assuming full penetrance when penetrance is in fact below 1 will result in a downwards bias of the tract length estimate.*

The results show that when the penetrance *P* is not 1, penetrance results in a downwards bias in the tract length estimation that is essentially *P*. Since a MLE of the penetrance can be obtained directly from the data (Hardarson et al. 2023), it is easy to correct for the assumption of no penetrance. That is, if penetrance is inferred to be below 1 for the dataset from which the gene conversion counts were obtained, simply scale the tract length estimates by $1/P$ since this (ignoring higher powers of $P$) is the factor by which penetrance inflates the ratio of single to multi-SNV conversions (see Figure S1.2). Scaling by $1/P$ is an approximation since this only corrects for the increase in single SNV conversions relative to double SNV conversions. The accuracy of the correction is improved if more conversions (and thus higher power of $P$) are considered.

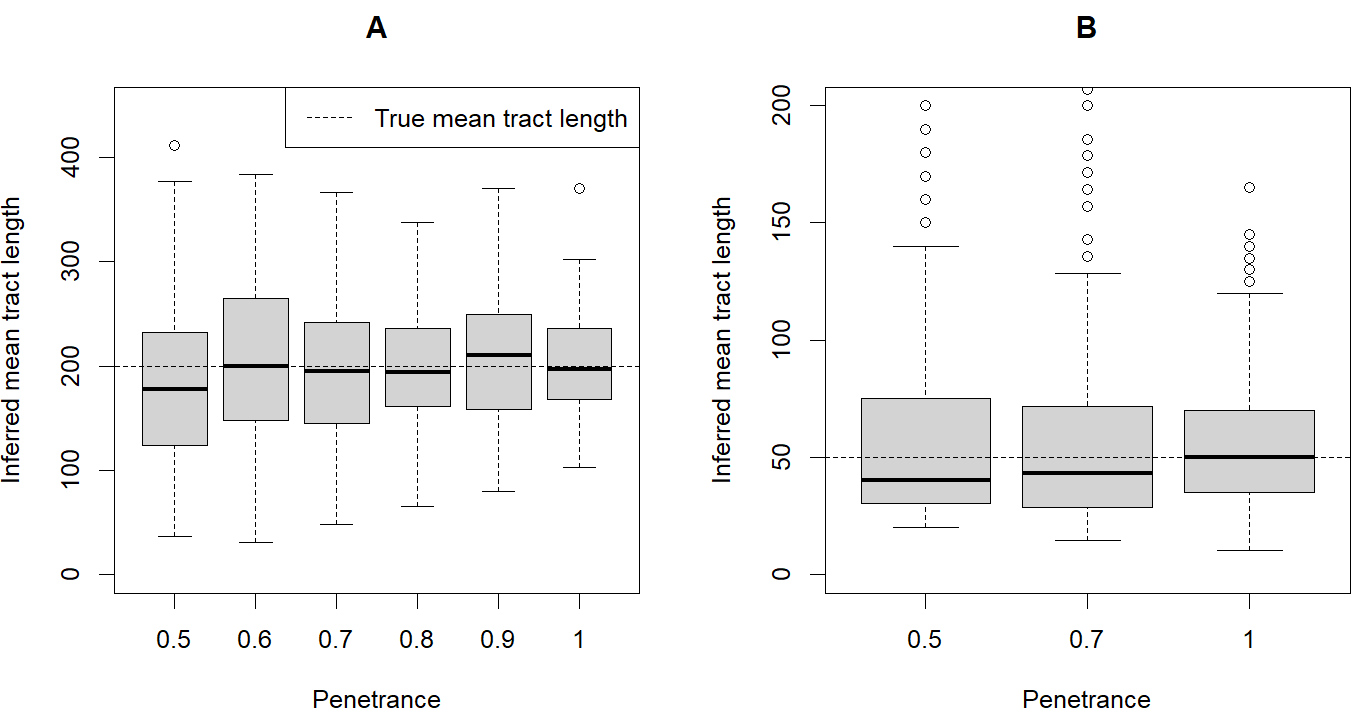

*Figure S1.2: Identical to Figure S1.1 but with all estimates multiplied by* $1/P$ *where P is the penetrance of the simulated dataset on which inference of the mean tract length was conducted. The results show that a model assuming full penetrance (*$P=1$*) can easily be applied to real data sets with* $P<1$ *since simply rescaling estimates by* $1/P$ *(to a first approximation) allows the correct tract length to be inferred even when* $P<1$*.*

**S2 - Read length effects on single/multi-snv conversion calls and detection**

Using our method to estimate NCO tract lengths with HiFi PacBio long read data could potentially result in an ascertainment bias since not all classes of gene conversion events (i.e. single, double, triple, etc) are equally likely to be distinguishable from crossover events. This is because the mean read length is 16.36 kb and calling a gene conversion event requires the converted SNV to be flanked by at least one unconverted SNV on both sides so as to distinguish the gene conversion event from a potential crossover event (see Porsborg et al. 2024 for details).

To test the effect of this potential ascertainment bias, we compared the tract length likelihood surfaces for some of our samples under conditions where gene conversions were called from simulated genome without read length limit to the case where the read lengths followed the distribution of our sample. We found little effect on the MLE of mean tract length is caused by ignoring read length distributions. This is because the average read length is several orders of magnitude larger than the average gene conversion tract (Figure S2.1A). However, a nearly 2-fold effect on detection probability (1-S(p,s), see main text) since finite reads means that flanking SNVs will not always be present on both sides of the converted SNV(s) which means the SNV cannot be called (Figure S2.1B). For details on SNV calling, see Porsborg et al. (2024). Because of this, the read distribution was taken into account when estimating detection probability and NCO rate (see main text, Figure 3).

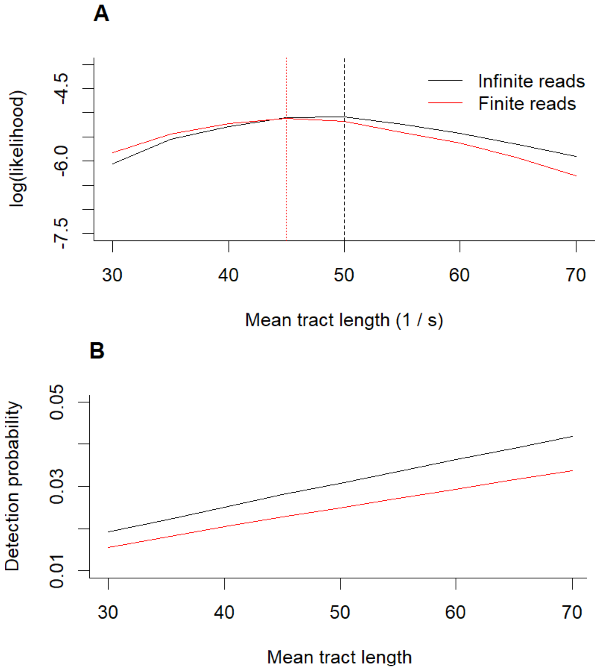

*Figure S2.1: Likelihood profiles obtained (at resolution of increments of 5 in 1/s) when simulating under infinite and finite and the actual read distribution are very similar. (A) Part of the likelihood surface for the sample where the likelihood profile is created by assuming unlimited read length and read lengths following the empirical read length distribution of the sample. Vertical dashed lines show the MLE given unlimited read length (black) and a read length distribution following the sample (red). (B) Detection probability against mean tract length under the assumption of unlimited read length (black) and read lengths following the empirical read length distribution of the sample (red).*

**S3 - The necessity of accounting for non-uniform SNV distributions**

We tested whether using the idealised model (which assumes a uniform distribution of SNVs) results in accurate inference of the mean tract length when using gene conversion counts from real genomic data (with a non-uniform SNV distribution). We did this by simulating gene conversion events with a known mean tract length throughout the genome of our sample individual (see methods). We then calculated the ratio of single to multi-SNV conversion events (Figure S3.1A, red) and compared to the ratio expected given a uniform SNV density (Figure S3.1.A, black).

We found that the idealised model overestimates the ratio under these conditions, since multi-SNV conversions are, for a given SNV density, far more likely in real genomes compared to a genome with a uniform SNV distribution. This is because some regions of the genome have very high SNV density (due to the coalescent, difference in mutation and recombination rate across the genome, and other processes). As a result, the idealised model also overestimates the mean tract length when the assumption of a uniform SNV distribution is violated (Figure S3.1. B). Because of this, we extend the model to obtain conversion probabilities given the exact SNV distribution of the sample, such that accurate estimates can be obtained even when the assumption of a uniform SNV distribution is completely violated (see main text).

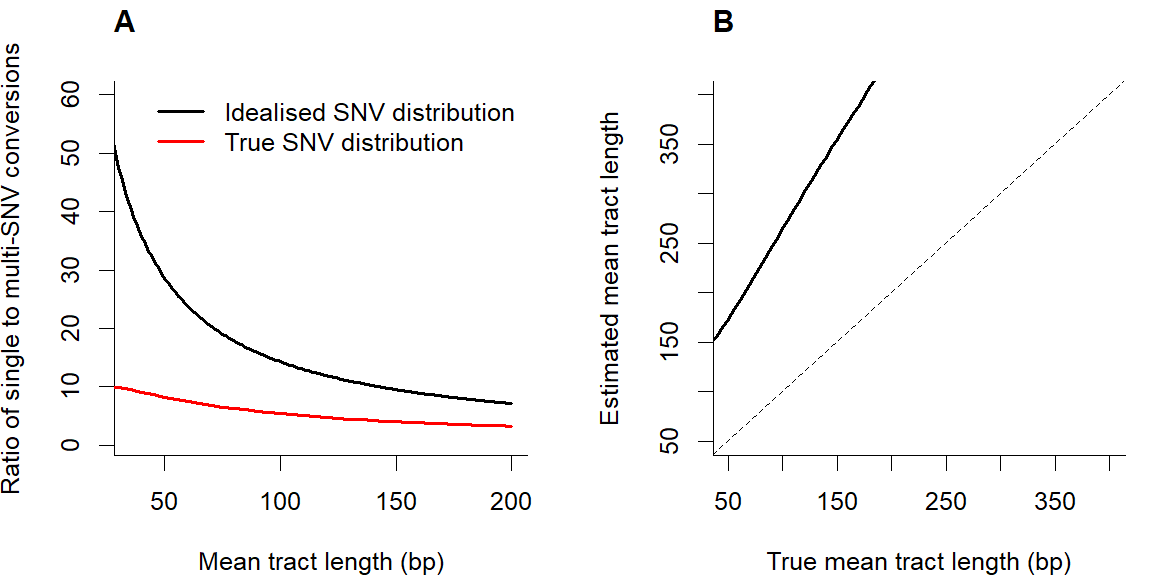

Figure S3.1. Evaluation of the idealised model when applied to genomic data with a non-uniform SNV distribution. (A) The idealised model predicts a much higher ratio of single to multi-SNV conversion than is actually observed (red). Because of this (B), the idealised model overestimates the mean tract length when the assumption of a uniform SNV distribution is violated. Dashed line shows x = y, i.e. perfect accuracy.

**S4 – model fit to gene conversion counts**

We tested whether the gene conversion counts (the number of single, double, triple, …etc gene conversion events) observed in our sample was compatible with tract being sampled from a single geometric distribution.

Specifically, we took the gene conversion counts and tested whether the counts obtained were significantly different from those expected if the counts were resulting from tracts sampled from a geometric distribution. That is, using the simulation approach outlined (see methods) we compared the probabilities of single, double, triple etc gene conversions under a geometric distribution with the numbers single, double, triple etc gene conversions observed in our sample. In our sample, we observed 167 single, 12 double 2, triple and 1 quadruple conversion events. Under the MLE s of 1/46 and given the SNV distribution of our sample, the expected counts of conversion events under a geometric distribution of tract lengths were 165.55 single, 13.20 double, 1.71 triple and 0.35 quadruple events.

The observed and expected counts were compared using a $\chi^{2}$-test, yielding $p=0.71$, suggesting that the gene conversion counts are not significantly different from what would be expected from a geometric distribution with the inferred MLE.

**S5 - SNV density at double-strand breaks**

Recombination (both crossover and NCOs) are thought to be initiated when a double-strand break (DSB) is catalysed by SPO11 which is recruited by PRDM9-binding sites throughout the genome (Baudat et al. 2010). Thus, NCOs are thought to be more likely to occur close to active PRDM9-motifs. However, if these motifs correlate (positively or negatively) with the SNVs density, this could bias our estimates of conversion probabilities and affect the accuracy of the estimates we present. To test this, we used the human recombination initiation map of Pratto et al. (2014) to test whether positions in the genome where DSBs (and hence NCOs) are likely, are correlated positively or negatively with the SNV density.

Following Pratto et al. (2014), while calling SNV and filtering reads via the pipeline given by Porsborg et al. (2024), we lifted the coordinates of DSB hotspots to the T2T reference genome (Nurk et al. 2022) and kept sequences for the hotspots seen in the AA-PRDM9 genotype which (given the European ancestry of our sample genome) is presumed to be the relevant PRDM9 genotype. This resulted in 35261 hotspot sequences, and these were BLASTed (Altschul et al. 1990) against the de novo assembly (see Porsborg et al. 2024 for details) to find the most likely positions of the hotspots within our sample genome. Centred at these 35261 positions, we counted the number of SNVs in a +10kb and -10kb window (i.e. 20kb window). We then randomly sampled 35261 20kb windows across the genome of our sample for comparison. We found that the SNV density differs by ~3% and the overall SNV distribution around DSBs is practically identical to the SNV distribution at randomly sampled positions across the genome (Figure S5.1). Given the robustness of our model, this ~3% difference in SNV density is unlikely to affect the estimate we present (See Figure 4, main text).

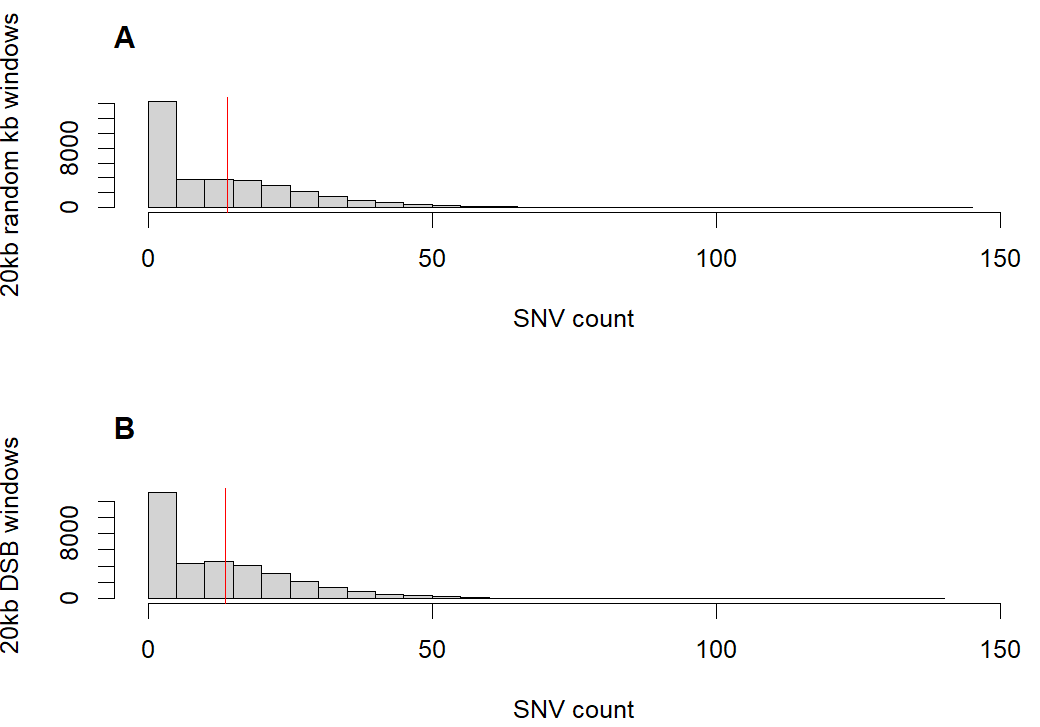

Figure S5.1: Distribution of SNV counts in 35261 windows of 20kb from (A) randomly sampled positions throughout the sample genome and (B) all DSBs found in our sample genome. Read lines show the average number of SNVs in a 20kb window and these are highly similar (0.0006997 SNVs/base in the genome in general and 0.0006790 SNvs/base around DSBs). CORR = -0.0012, P = 0.8225.

**S6: Accuracy and inference ability given sample size**

The idealised ratio model requires that both single and multi SNV tracts are observed. When tract length is very short, inference requires a lot of data because single SNV tracts become far more common than multi SNV tracts. We tested the extent of this effect by calculating the probability of observing at least one multi SNV tract given a certain number of observed tracts and some mean tract length (Fig S6.1). This can be interpreted as the probability of getting enough data for inference given sample size and mean tract length.

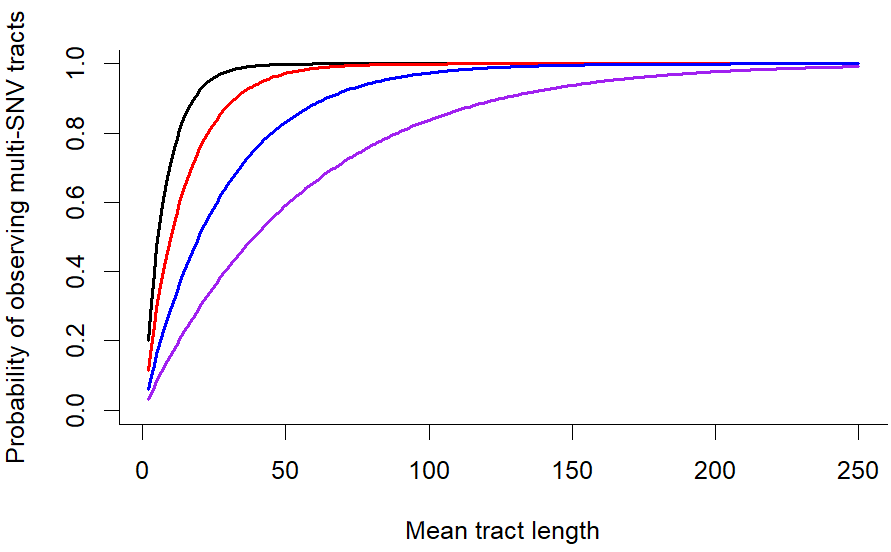

*Figure S6.1: The probability of observing at least one multi-SNV tract as a function of mean tract length given some total number of sampled tracts. Curves are shown for 182 samples (black) which was the number of observed tracts used for inference in this study (see main text) and 100 samples (red) 50 samples (blue) and 25 samples (purple). The calculations assume a uniform distribution of SNV and human density of SNVs (here assumed to be 0.00083 SNVs/base, see main text).*

**S7: Comparison of MLE to previous studies and methods**

To compare the estimate of mean tract length obtained, we compared the estimate directly to other estimates from previous studies. We found that while there is considerable variation (likely due to difference in inference methods, choice of distribution fitted to observed tracts, sequencing methods and choice of sample), our estimate agrees well with previous from humans overall (Figure S7.1) and estimate obtained from another mammal, namely from mice (Figure S7.2).

We note that this list of estimates is in no way exhaustive: we focus on recent studies (most published within the last decade) and we exclude studies which do not report at mean tract length or a range for the mean. These estimates should therefore be seen as examples of comparison between estimates, rather than an exhaustive list of all estimates within the literature. For a broader comparison of other estimates, especially from older studies, Mansai et al. 2011 gives a review. Further, it should be noted that there is considerable difference in how estimates are reported, making direct comparison more difficult (see legend; Figure S7.1-S7.2).

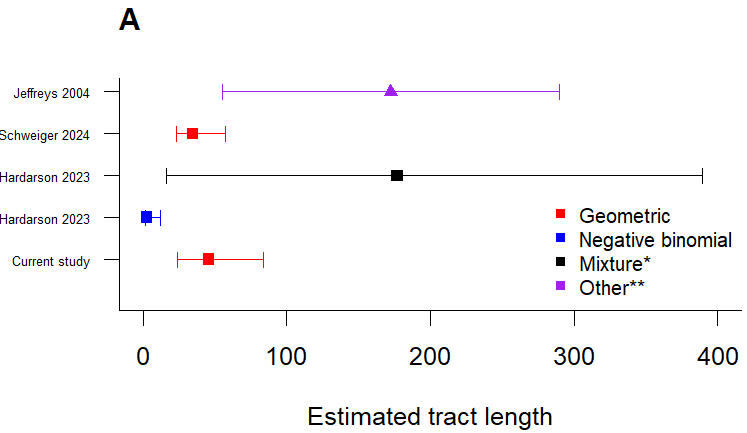

Figure S7.1: Tract length estimates from previous studies on male humans. Colours show the distribution of tract lengths fitted to the data. Squares show studies which report a mean with a 95% confidence interval and triangles show studies which report a different range containing a mean value. Hardarson et al. 2023 used data from Halldorsson et al. 2016.

*Mixture of two negative binomial distributions.

**Upper and lower bounds of the mean compatible with data found via simulation (see Jeffreys & May 2004 for details).

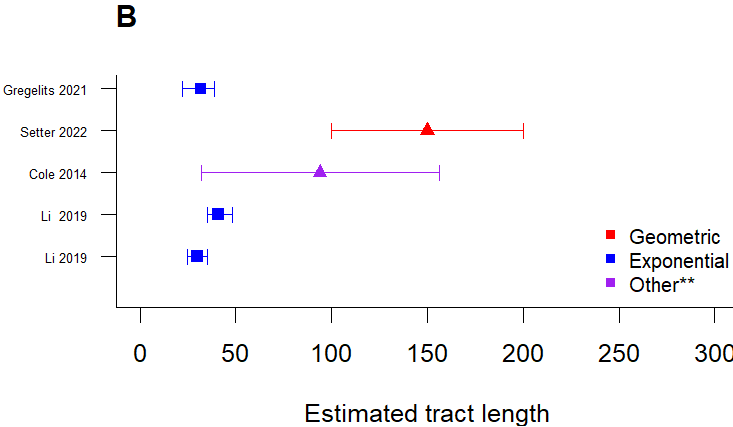

Figure S.7.2: Similar to S7.1, but with studies showing tract length estimates obtained from mice. Li et al. 2019 reports two means from different PRDM9 genotypes.

**Cole et al. 2014 used the midpoint method (see Mansai et al. 2011 for details).

To obtain a more direct comparison of the models, we applied the methods of Li et al. (2019) and Hardarson et al. (2023) to our data (Figure S7.3). We find that while some of the confidence intervals of the estimates obtained from the different methods overlap, the different methods yield differing estimates given the same data. This is due to the fact that the source of the information in the different methods are different. We use single/multi-conversions, Li et al. (2019) uses the distance between SNVs in the same gene conversion tract, and Hardarson et al. (2023) also uses information from the distance to flanking (unconverted) SNVs.

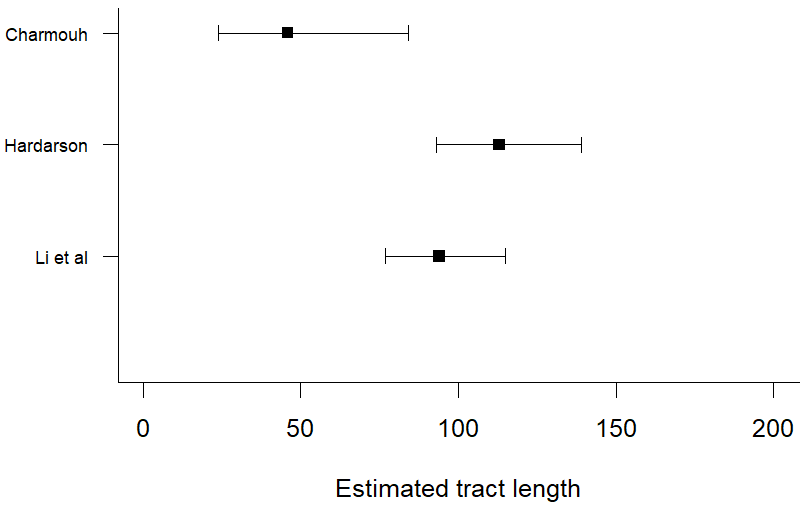

Figure S7.3: Estimates and confidence intervals for the mean tract length, using the methods described here as well as the methods described in (Hardarson et al. 2023) and (Li et al. 2019).
*To get the estimates using the methods described in (Hardarson et al. 2023) we assumed a geometric distribution.

**S8 – Detectability of long NCO events in HiFi long reads data**

To test whether the small proportion of so-called long NCO events recently reported (Halldorsson et al. 2016b; Wall et al. 2022; Versoza et al. 2023) were detectable using HiFi PacBio long read sequencing data, we simulated gene conversions in across our sampled genome. Here, we applied the read length distribution for the sample (approximated as a truncated normal distribution) with a mean read length of 16.36 kb, a standard deviation of 3.98 kb and a minimum and maximum read length of 11.44 kb and 24.14 kb, respectively. Using this read length distribution, we sampled reads and checked how often a gene conversion event within the read could be called (how often at least one SNV was converted while having flanking unconverted SNVs on the read). The results show that these long NCO event, which seem to occur at a very low rate, have a much higher detection probability (per event) than the shorter NCO events (Figure S8.1).

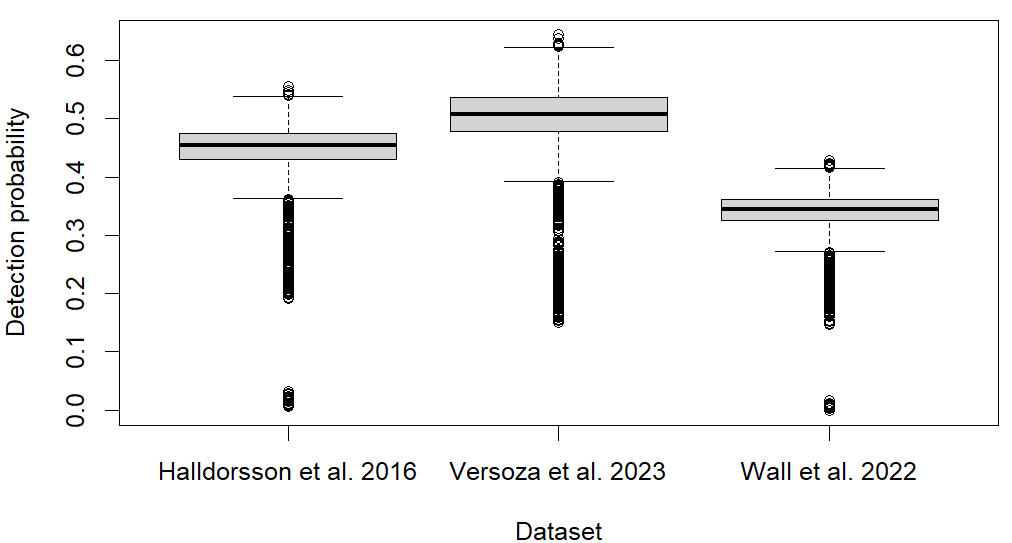
*Figure S8.1. Detection probability of long NCO events using mean long NCO lengths from different studies. Versoza et al. (2023) reported a mean of 5.166 kb, Wall et al. (2022) reported a mean of 47.58 kb and Halldorsson et al. (2016) reported a mode of 30kb (mean not reported). Using these reported means and modes as mean value of a geometric distribution, and assuming the SNV distribution of our sampled genome, the average detection probability of these long NCO events is about an order of magnitude higher than for the short NCO events. Each plot is based on 10^4^ simulations.*

**S9: Direct comparison between simulation procedure and recursion method**

Here we test whether the simulation approach to obtaining conversion probabilities ($P_{i}(s)$) results in the same conversion probabilities as the faster non-simulation approach (see Appendix) which relies on recursive calculation of tract counts and marginal tract counts. We test this for the probabilities of single, double, triple and quadruple conversions which are the types of conversions we find in the data (see Table 1, main text). We find that the simulation approach is in agreement with the recursive calculation approach for all these conversion probabilities when not accounting for read lengths (Figure S9.1).

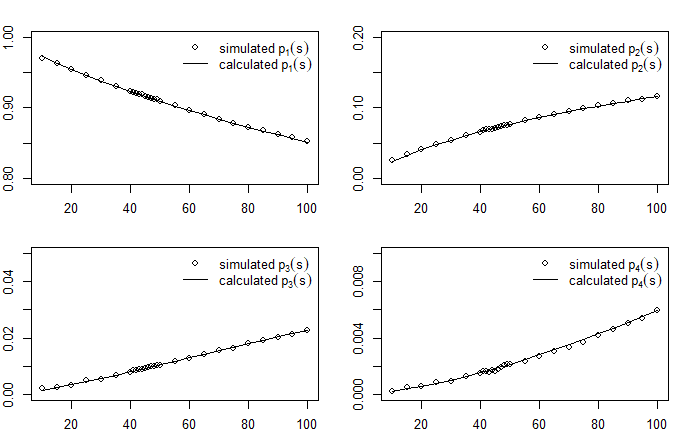

Figure S9.1: A comparison of $P_{i}(s)$ obtained via simulation (points) and recursive calculation (lines) for $i=1,2,3,4$ against $1/s$ (mean tract length). The results show that the recursive calculation of conversion probabilities is in agreement with simulation. Results shows assuming infinite read lengths.

Similarly, we find that the simulation approach is also in agreement with the recursive calculation approach for all these conversion probabilities when accounting for read lengths distribution of our sample (Figure S9.2).

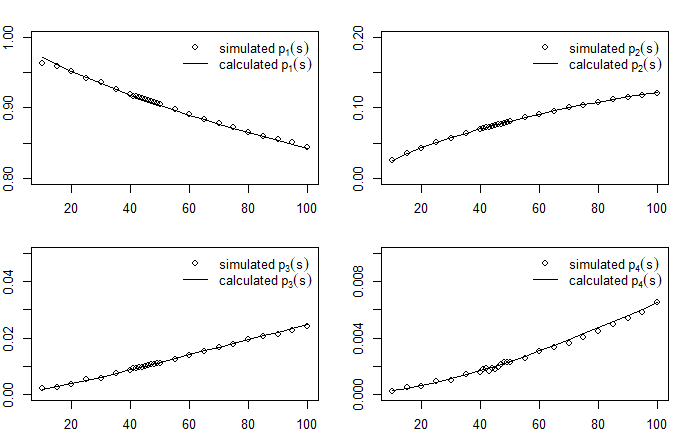

Figure S9.2: A comparison of $P_{i}(s)$ obtained via simulation (points) and recursive calculation (lines) for $i=1,2,3,4$ against $1/s$ (mean tract length). The results show that the recursive calculation of conversion probabilities is in agreement with simulation. Results shown for when the read length distribution of the sample is accounted for.
